# SerpinB3 drives cancer stem cell survival in glioblastoma

**DOI:** 10.1101/2021.12.21.473663

**Authors:** Adam Lauko, Josephine Volovetz, Soumya M. Turaga, Defne Bayik, Daniel J. Silver, Kelly Mitchell, Erin E. Mulkearns-Hubert, Dionysios C. Watson, Kiran Desai, Manav Midha, Jing Hao, Kathleen McCortney, Alicia Steffens, Ulhas Naik, Manmeet S. Ahluwalia, Shideng Bao, Craig Horbinski, Jennifer Yu, Justin D. Lathia

## Abstract

Despite therapeutic interventions for glioblastoma (GBM), cancer stem cells (CSCs) drive recurrence. The precise mechanisms underlying CSC therapeutic resistance, namely inhibition of cell death, are unclear. We built on previous observations that the high cell surface expression of junctional adhesion molecule-A drives CSC maintenance and identified downstream signaling networks, including the cysteine protease inhibitor SerpinB3. Using genetic depletion approaches, we found that SerpinB3 is necessary for CSC maintenance, survival, and tumor growth, as well as CSC pathway activation. The knockdown of SerpinB3 also increased apoptosis and susceptibility to radiation therapy. Mechanistically, SerpinB3 was essential to buffer cathepsin L-mediated cell death, which was enhanced with radiation. Finally, we found that SerpinB3 knockdown dramatically increased the efficacy of radiation in pre-clinical models. Taken together, our findings identify a novel GBM CSC-specific survival mechanism involving a previously uninvestigated cysteine protease inhibitor, SerpinB3, and provide a potential target to improve the efficacy of standard-of-care GBM therapies against therapeutically resistant CSCs.

**Summary:** Lauko et al. demonstrate a functional role for SerpinB3, which is elevated in glioblastoma cancer stem cells and protects against lysosomal-mediated cell death. SerpinB3 can be targeted to increase the efficacy of radiation in glioblastoma pre-clinical models.

## Introduction

Glioblastoma (GBM, WHO grade 4 glioma) is the most common primary malignant brain tumor and remains uniformly lethal. Despite aggressive therapies including maximal safe surgical resection, radiation, and chemotherapy, GBM patients experience a median survival of approximately 20 months (Stupp et al., 2017, 2005). GBM therapeutic resistance has been associated with poor brain penetration of compounds due to the blood-brain barrier (Bellettato and Scarpa, 2018; Harder et al., 2018), cellular heterogeneity and plasticity (Lauko et al., 2021), and limited immune infiltration (Martinez-Lage et al., 2019; Pombo Antunes et al., 2020). The cellular heterogeneity is driven by populations of cancer stem cells (CSCs) (Gimple et al., 2019; Lathia et al., 2015), and recent studies have demonstrated that GBM contains a high degree of plasticity, with the CSC state being linked to cellular programs including wound healing, development, and metabolic fluidity (Garnier et al., 2019; Mitchell et al., 2021; Pelaz et al., 2020) that underlie tumor growth and therapeutic resistance.

CSCs are functionally defined by their ability to self-renew and initiate a tumor upon secondary transplantation. Moreover, CSCs possess enhanced molecular mechanisms of therapeutic resistance, including DNA repair (Bao et al., 2006a) and drug efflux pumps. In addition, CSCs can be subject to stressful environments, including hypoxia and necrosis, and, although some mechanisms have been proposed as to how these cells can thrive under these stressful conditions (Alvarado et al., 2017; Hsieh et al., 2011), the precise mechanisms as to how CSCs evade cell death are unclear. Cell adhesion represents a cellular mechanism that promotes pro-survival signaling and is enhanced in CSCs (Lathia et al., 2014, 2010). Specifically, GBM CSCs present elevated expression of integrins (Lathia et al., 2010), cadherins (Siebzehnrubl et al., 2013) and junctional adhesion molecule-A (JAM-A) (Alvarado et al., 2016; Lathia et al., 2014), that drive self-renewal and promotes resistance to conventional therapies (Bao et al., 2006b; Colak and Medema, 2014). However, the CSC-specific intracellular signaling networks that link adhesion to resistance of cell death remain poorly defined.

Cell death is a complex and tightly regulated series of cellular programs that cancer cells have evolved to evade (Castelli et al., 2021; Safa, 2016). Cell death can be triggered through a variety of mechanisms including DNA damage, extrinsic ligands, and stressors (including hypoxic, metabolic, and endoplasmic reticulum stress). An understudied trigger of cell death is increased lysosomal permeability that leads to the release of reactive oxygen species and cathepsins (Wang et al., 2018), a diverse family of proteases (Yadati et al., 2020) that can initiate a variety of cell death programs. Under physiologic conditions, cathepsin proteolytic activity is buffered by protease inhibitors, including a family of serine(cysteine)-protease inhibitors termed serpins (Heit et al., 2013). In normal physiology, cathepsins and serpins exist in equilibrium to prevent aberrant cell death and damage to healthy tissues (Heit et al., 2013; Strnad et al., 2020). In GBM, there is limited information on lysosome-mediated cell death, which led us to hypothesize that CSCs might employ serpins to counteract this mechanism of cells death. Here, we show that the CSC programs regulated by JAM-A engages SerpinB3 downstream to simultaneously maintain the CSC phenotype and inhibit lysosome-mediated cell death. Suppression of SerpinB3 increases cell death, decreases self-renewal and tumor initiation, and enhances the response of CSCs to radiation via lysosomal-mediated cell death.

## Results

### SerpinB3 complexes with JAM-A and promotes the cancer stem cell phenotype in GBM

A series of cell surface receptors have been identified including CD133 (Liu et al., 2006; Singh et al., 2004), CD15 (Son et al., 2009), CD49f (Lathia et al., 2010), CD44 (Beier et al., 2007), L1CAM (Bao et al., 2008), and JAM-A (Lathia et al., 2014) that regulate the expression of the CSC phenotype. With rare exception, the molecular cascades that connect these receptors to the downstream pluripotency machinery, has not been made clear. In previous work, we established that GBM tumor cells expressed JAM-A when cultured under CSC-enriching conditions and that JAM-A expression was both necessary and sufficient for in vitro self-renewal. Subsequently, we revealed that Akt-activation functions downstream of JAM-A and could be inhibited by microRNA-145 (miR-145) (Alvarado et al., 2016). In total, we have assembled a putative signaling axis in which JAM-A regulates expression of the pluripotency machinery in GBM through Akt activation, however, this molecular cascade is far from complete.

To expand our understanding of JAM-A signaling, we sought out additional binding partners utilizing a histidine (His)-tagged JAM-A that we introduced into the T4121 GBM patient-derived xenograft model (PDX). We then pulled down the His-tagged JAM-A and identified a series of new binding partners using liquid chromatography coupled to mass spectrometry (**Fig. 1A, Supplemental Table 1**). As expected, the protein with the greatest number of peptides identified was JAM-A itself, and we narrowed down candidate hits from the initial list removing those with reported non-specific binding (via CRAPome (Mellacheruvu et al., 2013)). Post data filtration, our next strongest hit was a cysteine-protease inhibitor, SerpinB3 (also known as squamous cell carcinoma antigen 1 (SCCA1)), which has known roles in tumor progression in cervical, head and neck cancers, and hepatocellular carcinoma (HCC) (Cannito et al., 2015; Pontisso, 2014). We first validated binding of SerpinB3 to JAM-A (**Fig. 1B**). Further, SerpinB3 appears to be downstream of JAM-A, as reduction of JAM-A in cultured GBM tumor cells resulted in reduction of SerpinB3 (**Supplemental Fig. 1A**). We observed SerpinB3 expression in JAM-A-positive CSCs in a PDX model (**Fig. 1C**), as well as in human GBM patient tissue (**Supplemental Fig. 1B, C**). Transcriptional profiling indicated that SerpinB3 expression was not specifically associated with any given molecular subtype (data not shown). Together, these data indicate the SerpinB3 expressed under CSC-optimized conditions in vitro and heterogeneously in CSCs and GBM patient tumor specimens.

**Figure 1.**
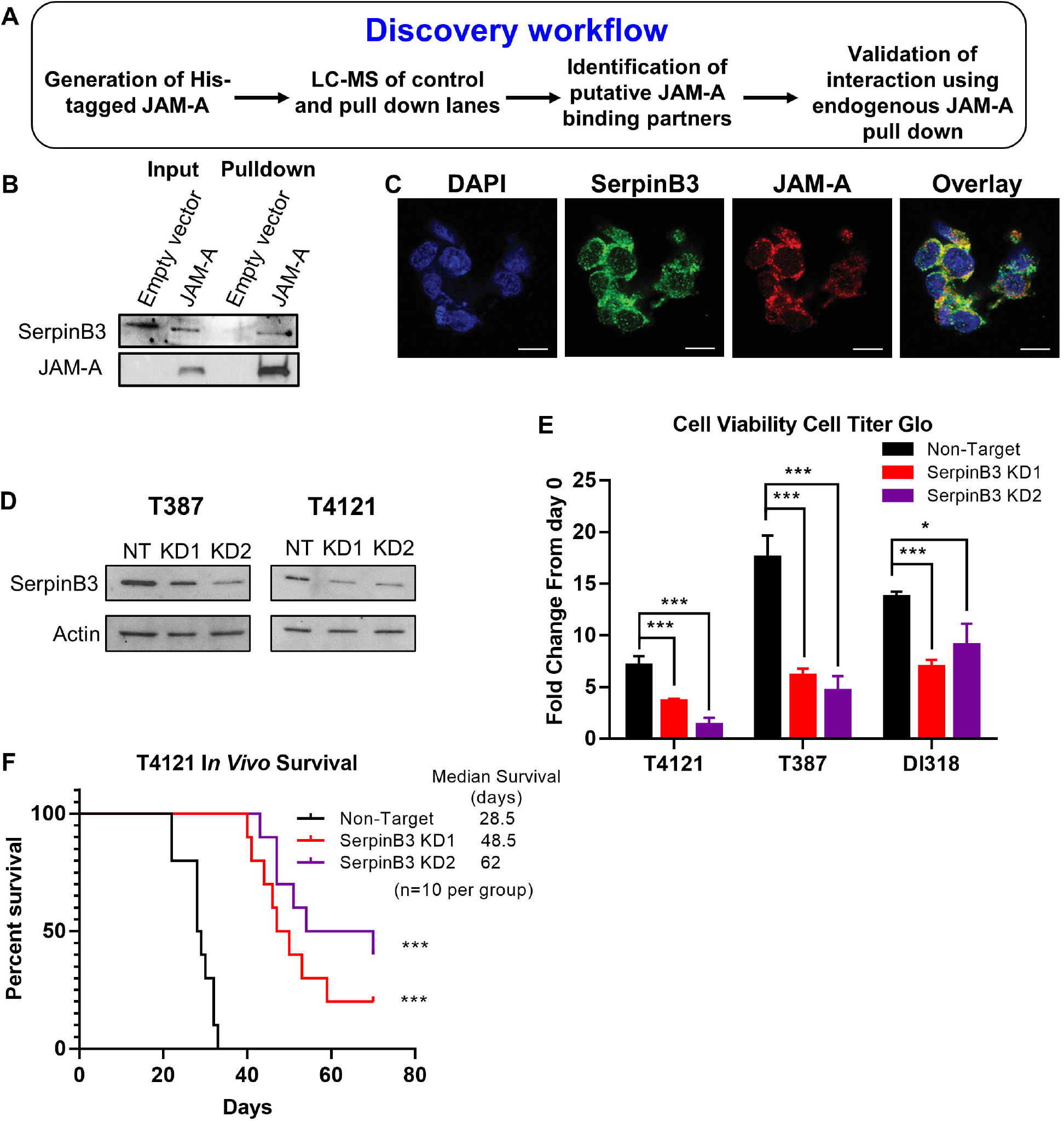
SerpinB3 is necessary for glioblastoma. A) Graphical abstract of His-JAM-A pulldown and LC-MS procedure. B) Verification of LC-MS results by western blot. C) Immunofluorescent staining demonstrating co-expression of JAM-A and SerpinB3 in T387 PDX glioblastoma tumor model (scale bar 10 μm). D) Western blot demonstrating knockdown of SerpinB3 with each shRNA, KD1 and KD2. Actin was utilized as a loading control. E) Fold change in cell viability at day 7, normalized to day 0, in three PDX glioblastoma models. Cell viability measured with CellTiter-Glo Luminescent Cell Viability Assay (5 technical replicates per condition, per tumor model). F) Kaplan-Meier curves depicting survival of mice with 20,000 T4121 tumor cells intracranially injected. Cells were transfected with either non-target (SHC002) or SerpinB3 (KD1 or KD2) shRNA, with n=10 mice per group. p<0.05 was considered statistically significant. *, p<0.05; **, p<0.01; ***, p<0.001 as determined by one-way ANOVA with Dunnett’s multiple comparisons test or log-rank test for survival data. Error bars represent standard deviation.

Given the limited understanding of the role of SerpinB3 in GBM, we sought to assess its function using a genetic depletion approach. We utilized non-overlapping shRNAs against SerpinB3 and were able to reduce protein levels in multiple GBM PDX-derived CSC models (T4121, T387, DI318) compared to non-targeting (NT) controls (**Fig. 1D, Supplemental Fig 1E**). While foundational experiments were performed using four different shRNAs, we subsequently focused on two constructs, knockdown 1 (KD1) and knockdown 2 (KD2). The reduction of SerpinB3 in CSCs resulted in a decreased number of viable cells in vitro (**Fig. 1E, Supplemental Fig. 1D**,**F**) and a potent reduction in tumor initiation in vivo (**Fig. 1F**). Additionally, we utilized the Depmap portal (https://depmap.org/portal/interactive/) that combines RNAi screens from over 31 GBM cell lines to determine the dependency of these cells on SerpinB3. Of the 31 cell lines, all but four were dependent on SerpinB3 (**Supplemental Fig. 1G**). Taken together, these data provide evidence that SerpinB3 is essential for GBM CSC growth and tumor initiation.

Given the phenotypes observed upon SerpinB3 knockdown in CSCs, we also assessed changes in CSC maintenance as a result of SerpinB3 depletion. We observed that SerpinB3 knockdown reduced CSC signaling via changes in mRNA levels of core pluripotency transcription factors (*OCT4, NANOG, MYC*) and CSC transcription factors (*OLIG2*) (**Fig. 2A**). We then interrogated the functional consequences of SerpinB3 knockdown on self-renewal via in vitro limiting-dilution assays, a surrogate for self-renewal that also can also be impacted by cell proliferation and cell death, and found a potent reduction in self-renewal with SerpinB3 knockdown compared to NT control conditions across multiple CSC models (**Fig. 2B**). To gain further insight into SerpinB3-mediated changes, we focused on c-myc and transforming growth factor-beta 1 (TGF-β1) based on their reported roles in GBM CSCs (Anido et al., 2010; Bruna et al., 2007; Wang et al., 2008), essential role in cancer cell proliferation, and link to SerpinB3 in hepatocellular carcinoma (HCC) (Turato et al., 2015, 2014). As predicted, knockdown of SerpinB3 reduced c-myc expression (**Fig. 2C**) and TGF-β1 secretion (**Fig. 2D**). To gain additional mechanistic insight into the role of SerpinB3 in CSC-mediated cell growth and tumor initiation, we subjected SerpinB3-depleted CSCs to a cancer-focused mRNA panel using the NanoString platform. Using an unbiased clustering, we observed that NT control samples were distinct from SerpinB3 knockdown (using the KD2 construct) in 2 CSC models (T4121, DI318, **Supplemental Fig. 2A**). We found a series of pathways that were differentially expressed in NT control compared to SerpinB3-depleted cells, including cancer driver genes and a number of pathways known to regulate CSCs, including Hedgehog, Notch, and TGF-β (**Fig. 2E, Supplemental Fig. 2B-D**). This finding corroborated our observation of reduce TGF-β1 secretion with SerpinB3 knockdown (**Fig. 2D**) and we additionally validated that SerpinB3 depletion reduced the expression of members of the Notch signaling network (*NOTCH2, JAGGED2*, **Supplemental Fig. 2E**) after SerpinB3 knockdown. These data indicate that SerpinB3 is essential for self-renewal and interacts with multiple CSCs signaling network nodes.

**Figure 2.**
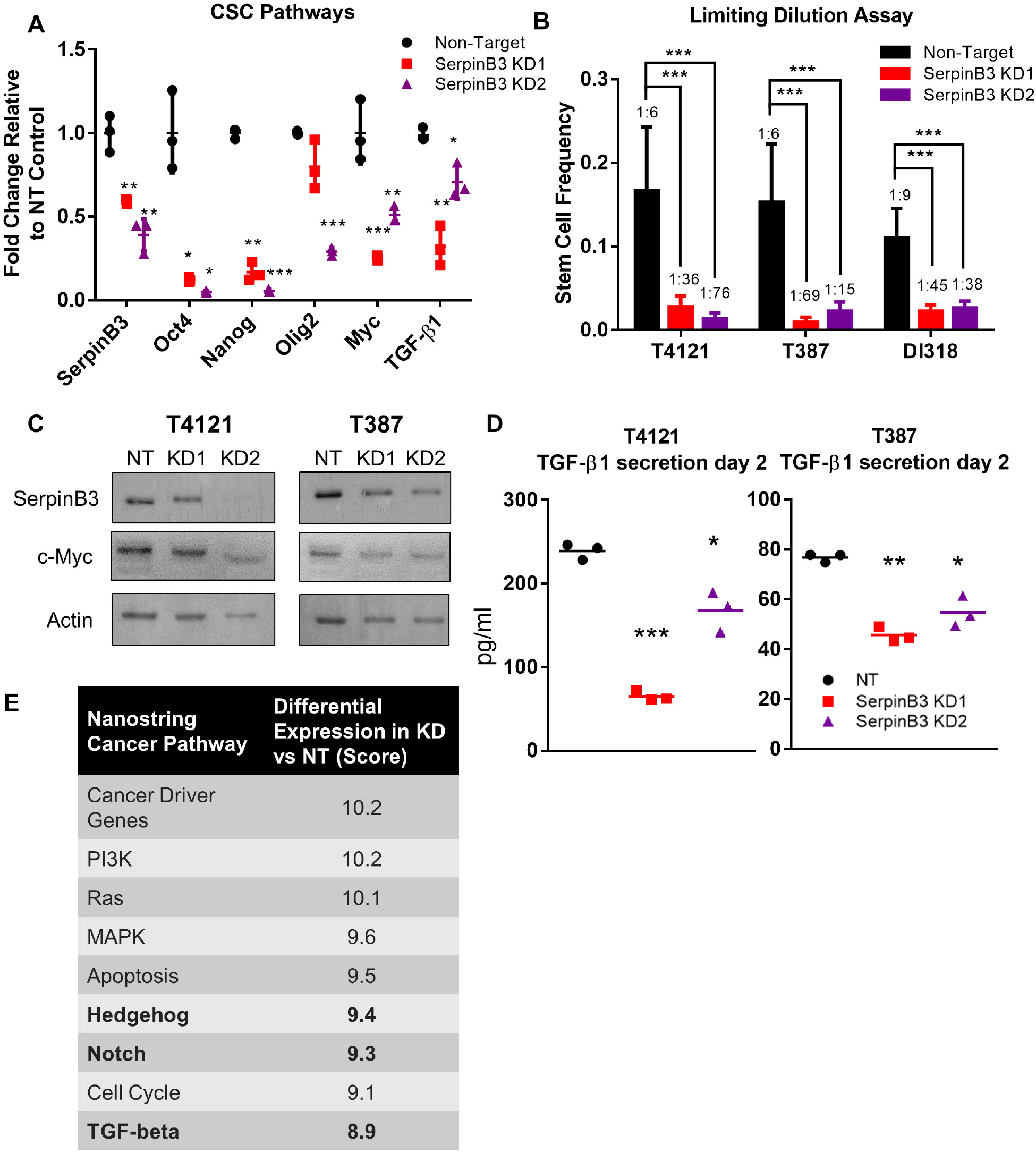
SerpinB3 regulates known CSC pathways. A) RNA was isolated after SerpinB3 knockdown, and qPCR was performed for *SERPINB3, OCT4, NANOG, OLIG2, MYC, and TGF-*β*1* (three technical replicates). B) Tumor cells were plated in a limiting-dilution manner, and the number of wells containing spheres was counted after 14 days and used to calculate stem cell frequencies using the online algorithm detailed in the methods. C) c-Myc expression after SerpinB3 knockdown was assessed via western blot, with actin as a loading control. D) TGF-β1 secretion was analyzed 2 days after plating and normalized to total protein (three technical replicates per condition, per tumor model). E) NanoString Pathway score comparing SerpinB3 knockdown to non-target control. Bolded rows represent pathways known to regulate the cancer stem cell state. p<0.05 was considered statistically significant. *, p<0.05; **, p<0.01; ***, p<0.001 as determined by one-way ANOVA with Dunnett’s multiple comparisons or chi-squared p-value for limiting dilution assay. Error bars represent standard deviation.

### SerpinB3 protects GBM tumor cells from apoptotic death

Based on the decrease in cell viability we observed after SerpinB3 knockdown and the well-established role for SerpinB3 in inhibiting cell death, we asked whether this correlates with an increase in cell death (Villano et al., 2014). Moreover, as our NanoString analyses of cancer pathways also revealed an increase in apoptosis with SerpinB3 knockdown compared to control conditions (**Fig. 2E)**, we validated this increase in apoptosis after SerpinB3 knockdown in multiple CSC models as read out by annexin V/PI double-positive cells (**Fig. 3A, B**). In addition, SerpinB3 knockdown also resulted in an increase in caspase 3/7 activity compared to control conditions in multiple CSCs models using the CaspaseGlo DEVD-aminoluciferin assay (**Fig. 3C, Supplemental Fig. 3A**). Additionally, we utilized the IncuCyte Caspase 3/7 assay, a DEVD-tagged DNA-intercalating dye, which also demonstrated increased caspase 3/7 activity (**Fig. 3D, E**). Combined, this data demonstrates an enhanced cell death phenotype after loss of SerpinB3.

**Figure 3.**
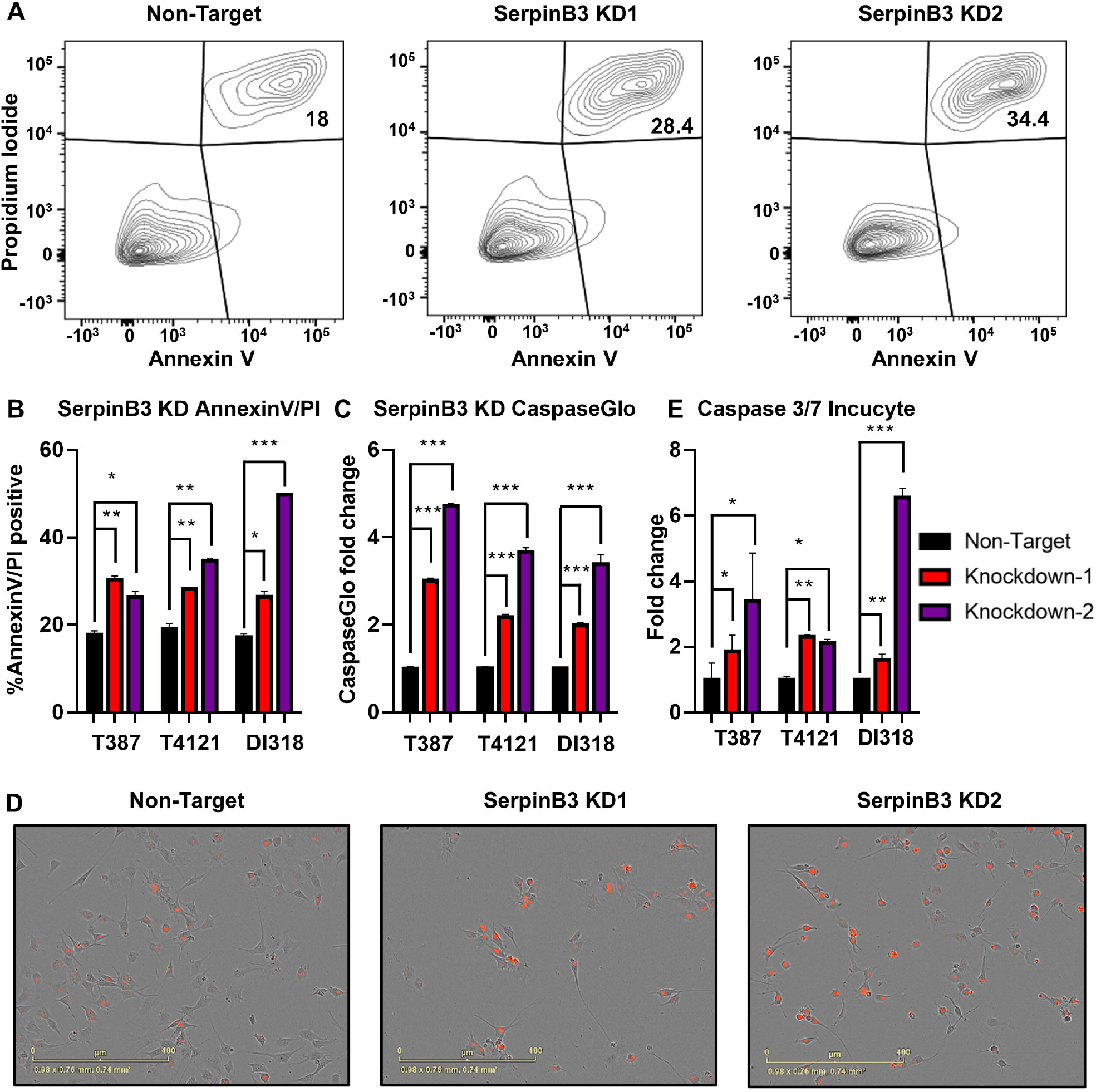
SerpinB3 inhibits cell death. A) Annexin V- and propidium iodide-positive sample flow plots (T4121). B) The percentage of annexin V- and propidium iodide-positive cells after SerpinB3 knockdown was compared to the non-target control group (three technical replicates per condition, per tumor model). C) Activity of caspase 3/7 was measured using Caspase-Glo after SerpinB3 knockdown, and the fold change compared to the non-target control is shown (three technical replicates per condition, per tumor model). D) Images from caspase 3/7 assay; red represents active caspase-positive cells (DI318). E) Quantification of caspase 3/7 IncuCyte assay normalized to cell confluence per well and compared to the NT condition (four technical replicates per condition, per tumor model). p<0.05 was considered statistically significant. *, p<0.05; **, p<0.01; ***, p<0.001 as determined by one-way ANOVA with Dunnett’s multiple comparisons. Error bars represent standard deviation.

### SerpinB3 inhibits lysosomal-mediated apoptosis

As apoptosis can be initiated via multiple pathways (i.e., intrinsic vs extrinsic), we sought to better understand the molecular mechanism though which SerpinB3 prevents cell death. SerpinB3 is a known inhibitor of cathepsin L, a cysteine protease that can relocalize to the cytoplasm after lysosomal membrane disruption and trigger lysosomal-mediated cell death (**Fig. 4A**) (Fehrenbacher et al., 2004; Oberle et al., 2010; Piazza et al., 2007). Initially, we used L-leucyl-L-leucine methyl ester (LLME) to compromise lysosomal membrane integrity and observed a potent decrease in cell viability, which was further enhanced with SerpinB3 knockdown (**Fig. 4B, Supplemental Fig. 3C**). We next investigated whether standard-of-care radiation treatment could impact lysosomal membrane integrity and found an increase in lysosomal membrane permeability 6 hours post-irradiation with a single dose of 5 Gy as read out by a shift in acridine orange localization (**Fig. 4C, D**), which fluoresces red in acidic compartments and green in the remainder of the cell. Additionally, we observed a relocalization of cathepsin L, a direct target of SerpinB3 (Sun et al., 2017), from the lysosome to the cytoplasm at 6 hours after irradiation with a single 5 Gy dose (**Fig. 4E, F; Supplemental Fig. 3B**). This can be observed as a shift in the cathepsin L from primarily puncta to a more diffuse localization throughout the cytoplasm. Importantly, we did not observe an increase in overall cathepsin L or SerpinB3 levels at this time point (**Supplemental Fig. 3D**). As radiation is part of standard of care for GBM and as CSCs are resistant to radiation, we next assessed whether depletion of SerpinB3 increases the efficacy of radiation in CSCs. SerpinB3 knockdown potently increased the sensitivity of CSCs to radiation compared to control NT conditions (**Fig. 4G**). Taken together, these data indicate that SerpinB3 protects against lysosomal membrane permeability induced by radiation.

**Figure 4.**
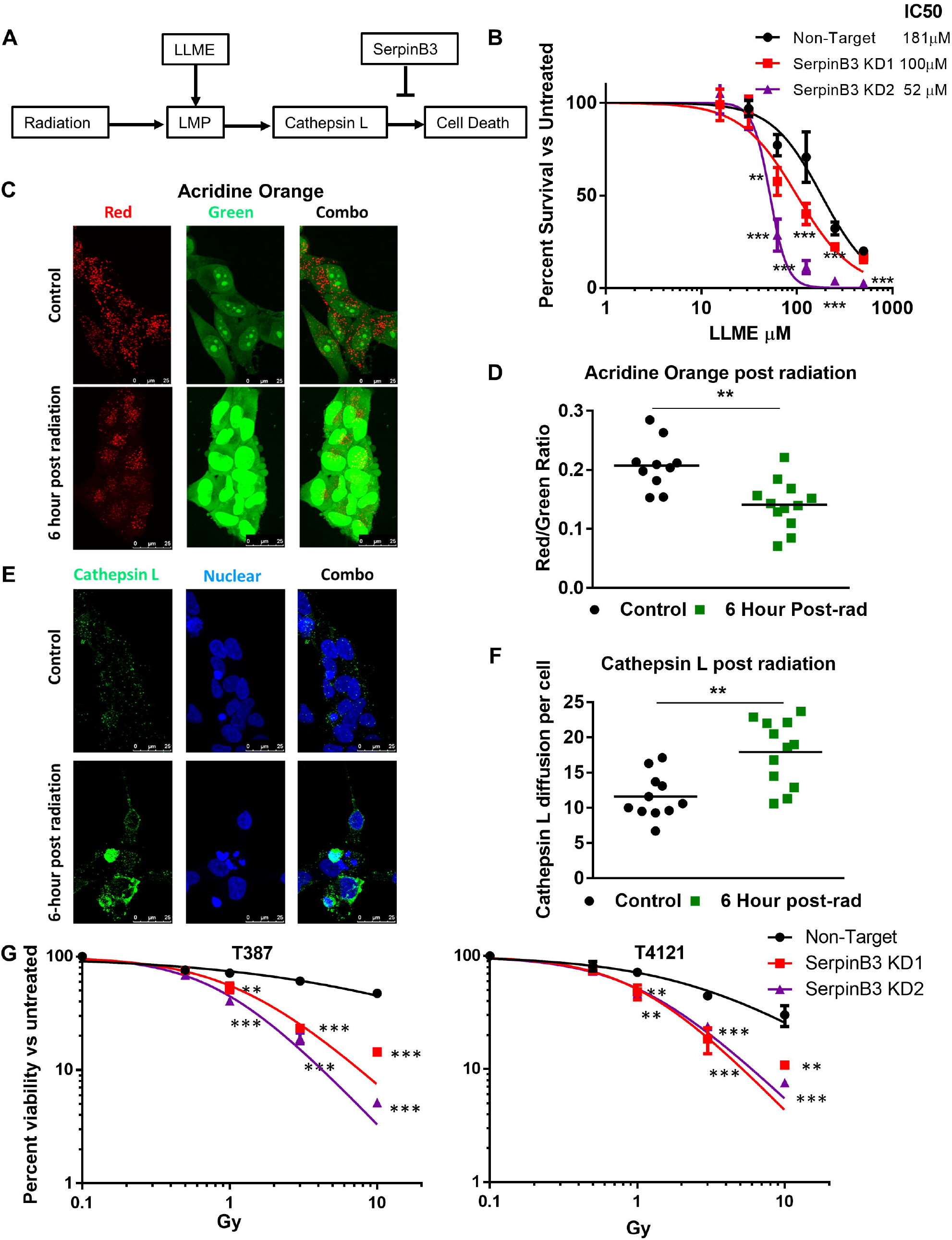
SerpinB3 buffers cells from lysosomal membrane permeability. A) Schematic of lysosomal mediated cell death after radiation B) T387 cells were treated with LLME at varying concentrations, and the IC50 was determined (five replicates per condition). C-D) Acridine orange was added to cells 6 hours after irradiation with 5 Gy, and images from 12 random visual fields were taken. The red/green ratio per image was calculated comparing control to irradiated conditions. E-F) Six hours post irradiation, cells were fixed with paraformaldehyde and stained for cathepsin L. The integrated density of cathepsin L per cell was determined and compared between the radiation and control conditions (12 images per condition). G) Cell viability was measured after two days of varying doses of radiation, and the percent of viable cells is shown compared to each group’s untreated control at each dose of radiation (three technical replicates per condition, per tumor model). p<0.05 was considered statistically significant. *, p<0.05; **, p<0.01; ***, p<0.001 as determined by one-way ANOVA with Dunnett’s multiple comparisons. Error bars represent standard deviation.

### SerpinB3 loss enhances radiation in vivo

To determine whether SerpinB3 is important for radiation resistance in vivo, we transplanted NT and SerpinB3-knockdown T4121 CSCs and subjected the mice to a pre-clinical radiation paradigm (**Fig. 5A**). We found that SerpinB3 knockdown increased tumor latency, and this was further extended by irradiation with 10 Gy over 5 days, increasing the hazard ratio in SerpinB3 knockdown compared to control conditions (**Fig. 5B-E**). This response was also observed with a lower dose of radiation (**Supplemental Fig. 4A-E**). Taken together, these data suggest that SerpinB3 prevents cell death and contributes to radiation resistance.

**Figure 5.**
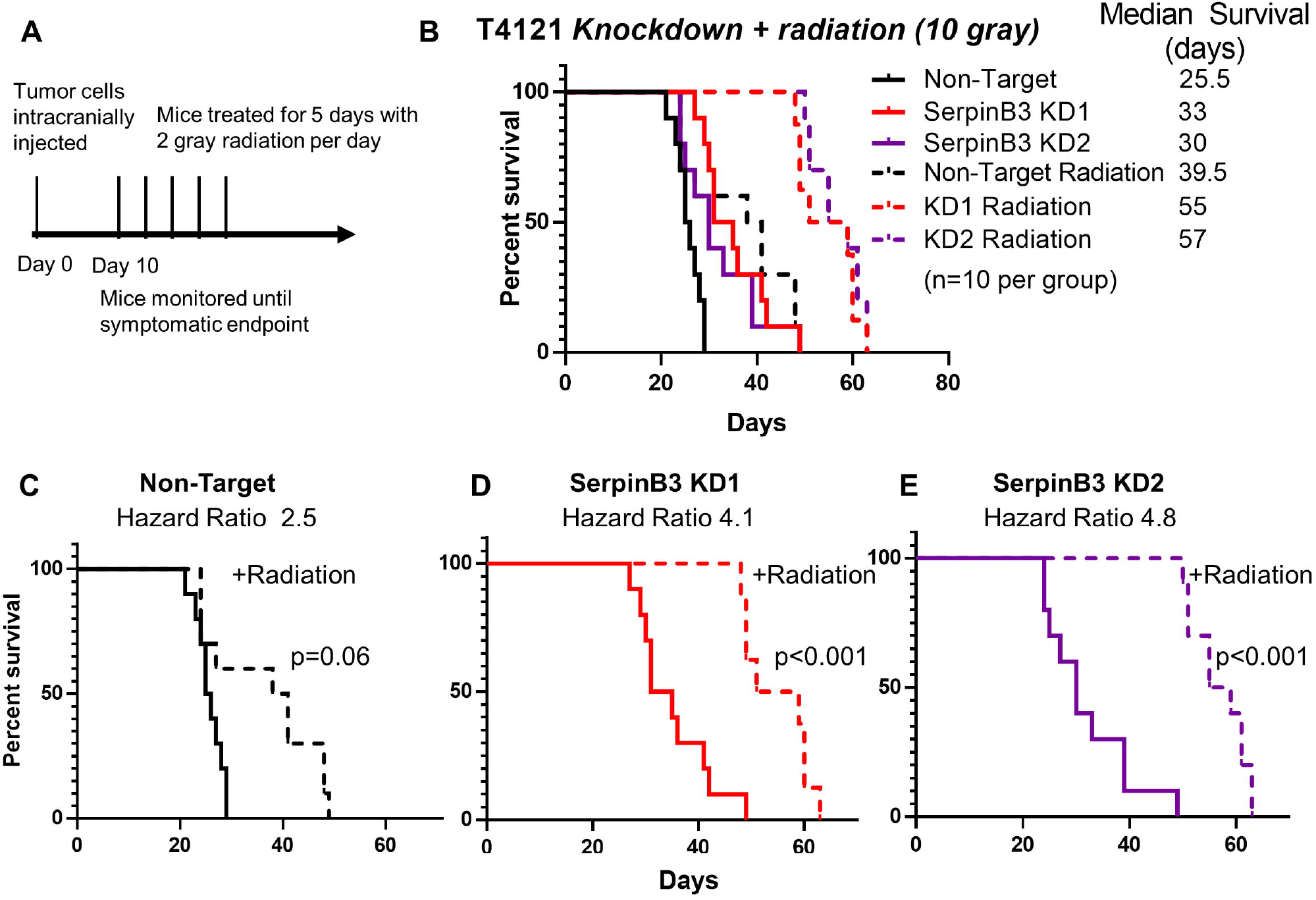
SerpinB3 contributes to radiation resistance. A) Schematic of in vivo radiation experiment with 10 Gy total radiation treatment. B-E) A total of 20,000 tumor cells per condition were intracranially injected into 10 mice per group. Ten days after irradiation, mice received 2 Gy of radiation per day for 5 days (total of 10 Gy) to the head. B) All treatment groups are shown together with median survival values given. The groups were subsequently divided into: C) Non-target with or without radiation, D) SerpinB3 KD 1 with or without radiation, and E) SerpinB3 KD 2 with or without radiation. * p<0.05 was considered statistically significant. *, p<0.05; **, p<0.01; ***, p<0.001 as determined by log-rank test.

## Discussion

Resistance to apoptosis is a well-recognized hallmark of cancer, but specific resistance mechanisms underlying cell death have not been thoroughly investigated in CSCs. This is surprising given the predominant phenotype of CSCs is enhanced therapeutic resistance, specifically to radiation and temozolomide in the case of GBM. SerpinB3 represents a new mechanism for GBM CSC survival that may also be functionally important in other cancers. The role of SerpinB3 in cancer is not well developed, despite being originally identified as overexpressed in squamous cell carcinoma (Kato and Torigoe, 1977). Studies in cervical cancer, non-small cell lung cancer, breast cancer, esophageal squamous cell carcinoma, and hepatocellular carcinoma have correlated elevated SerpinB3 expression with clinical stage and decreased response to therapy (Collie-Duguid et al., 2012; Liu et al., 2015; Ngan et al., 1990; Petty et al., 2006; Shimada et al., 2003). SerpinB3’s role in GBM had not been studied until recently when a novel lncRNA, TMEM44-AS1 (Bian et al., 2021), was found to bind to SerpinB3, forming a positive feedback loop with Myc. While this suggests an importance of SerpinB3 in GBM, the detailed molecular mechanism(s) by which SerpinB3 drives oncogenesis in its role in therapy resistance have yet to be fully elucidated.

There are several hypothesized mechanisms by which SerpinB3 could affect cancer-relevant phenotypes (impact on the stem cell state, resistance to apoptosis, invasion). A recent study in cholangiocarcinoma found that SerpinB3 was expressed in a stem-like subset of cells and that knockdown of SerpinB3 resulted in decreased invasion and proliferation (Correnti et al., 2021). In addition to its roles in cancer, SerpinB3 is expressed in hepatic stem cells, where SerpinB3 expression correlates with decreased activated caspase 3 (Villano et al., 2014). SerpinB3 has been linked to the inhibition of apoptosis in the settings of endoplasmic reticulum stress (Verfaillie et al., 2013), TNF-alpha release (Suminami et al., 2001), radiation (Murakami et al., 2001) and ultra-violet-radiation (Katagiri et al., 2006), but the exact mechanism unknown. In this study, we have highlighted the role of lysosomal membrane permeability and cathepsin L release as a mechanism of resistance to apoptosis. SerpinB3 inhibits cathepsin L (Sun et al., 2017), and cathepsin L released from the lysosomes and other acidic compartments has been shown to cause caspase-mediated cell death (Fehrenbacher et al., 2004; Oberle et al., 2010). Finally, while multiple mechanisms of radiation-induced cell death have been documented, our study builds on earlier work highlighting the role of lysosomal membrane permeability in sensitizing tumor cells to radiation in GBM (Zhou et al., 2020). Taken together, these data outline a novel mechanism whereby SerpinB3 expressed in GBM tumor cells leads to radiation resistance by buffering lysosomal membrane permeability.

Targeting the CSC-state remains a clinically interesting possibility and our observations suggest that SerpinB3 inhibition may be a mechanism by which cells expressing this phenotype can be sensitized to radiation. Our observations, the upregulated expression of SerpinB3 in other cancers as well as the dependency of other tumor cells on SerpinB3 (**Supplemental Fig. 4F)**, set the foundation for additional studies of stem-like cells in other tumor types. While this is one of the first examples of SerpinB3 function in GBM CSC populations, our studies have some limitations. Our observations were made using genetic approaches, and future priorities include development of brain-penetrant SerpinB3 inhibitors. The flexibility of the protease inhibitor domain represents one such therapeutic development challenge, along with the high degree of homology between SerpinB3 and SerpinB4 outside the protease inhibitor region. These challenges necessitate further understanding of this newly identified signaling network, including JAM-A dependent vs independent functions of SerpinB3 and the importance of JAM-A SerpinB3 stability, localization, and function. Another consideration for therapeutic development is the impact on cell death, which is a fundamental process in organ development and homeostasis in healthy tissue. The extent to which inhibition of SerpinB3 may serve as a transforming event for cancer initiation should be an additional consideration. Despite these limitations, SerpinB3 has emerged as a molecule of interest across numerous tumor types and further research into mechanisms for targeting SerpinB3 is required moving forward. In summary, our findings suggest that SerpinB3 may be a targetable mechanism leveraged by CSCs to resist cell death (**Fig 6)**.

**Figure 6.**
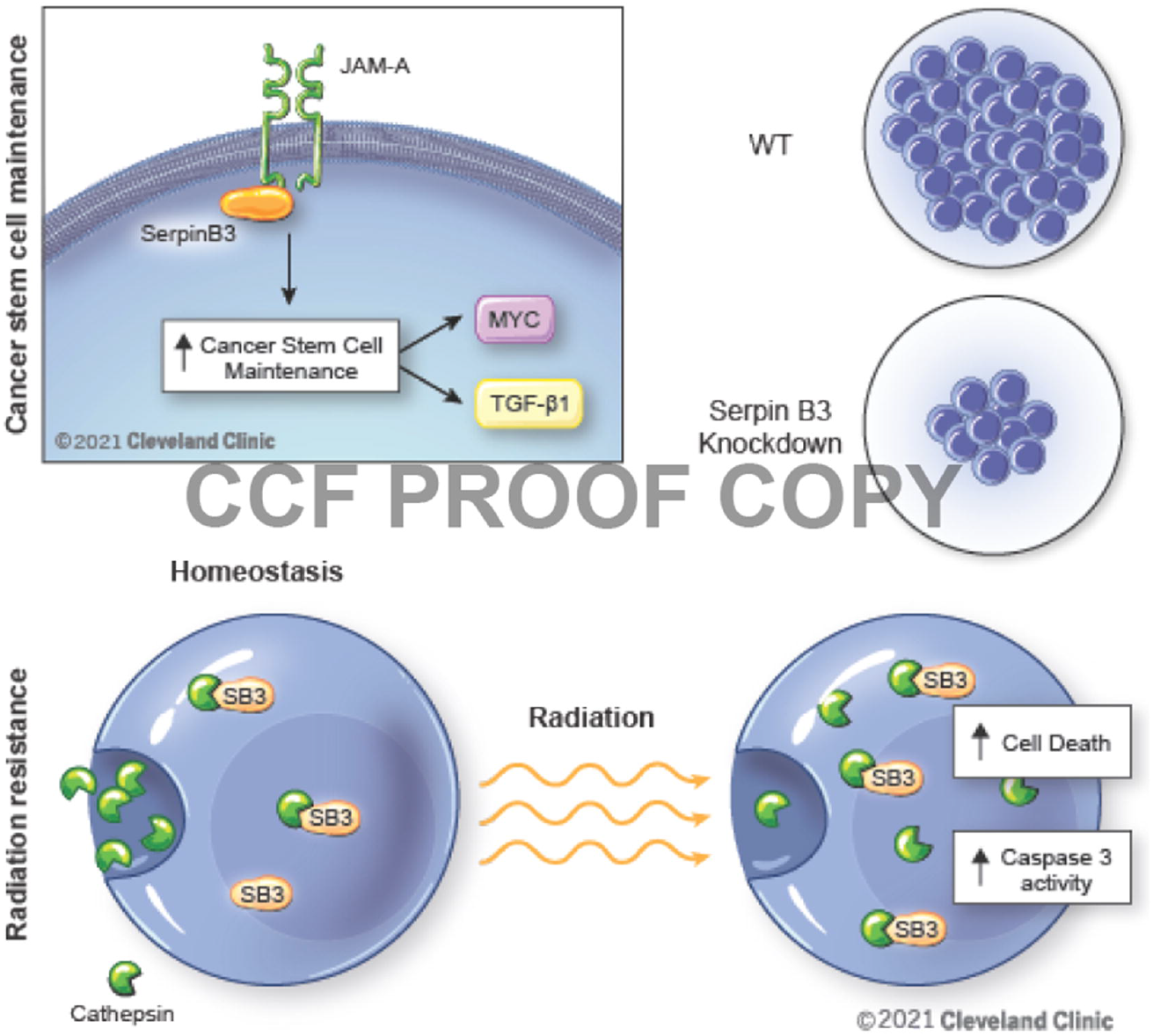
drives cancer stem cell survival in glioblastoma.

## Materials and Methods

### GBM Tumor cell derivation and culture

GBM tumor models were generated by passaging primary tumor cells through immunocompromised mice as previously described (Bao et al., 2006b; Lathia et al., 2010). Briefly, primary tumor cells were intracranially implanted into NOD.Cg-*Prkdc*^*scid*^ *Il2rg*^*tm1Wjl*^/SzJ (NSG) mice, and upon tumor formation, tumors were isolated and digested with papain (Worthington). Dissociated cells were plated overnight in Neurobasal™ Medium minus phenol red (ThermoFisher) with 1x B27 supplement (ThermoFisher), 1 mM sodium pyruvate, 2 mM L-glutamine, 50 U/mL penicillin/streptomycin, 20 ng/ml human (h)EGF and 20 ng/ml hFGF2 (R&D systems). Subsequently, CD133+ cells were isolated by magnetic bead sorting (Miltenyi) and cultured in the media described above. Some cell models were previously established at Duke University and obtained through approved material transfer agreements. CD133+ cells were seeded in suspension culture at 5×10^4^ cells/ml and passaged no more than 10 times. After 10 passages, cells were re-implanted into NSG mice and enriched for CD133+ cells. De-identified GBM specimens were collected from the Cleveland Clinic Brain Tumor and Neuro-Oncology Center in accordance with an Institutional Review Board-approved protocol, and informed consent was obtained from all GBM patients contributing tumor specimens.

### Immunoblotting

Protein was isolated from cells using a lysis buffer composed of 10 mM Tris HCl, 1 mM EDTA, 150 mM NaCl, 0.5% NP-40, 1 mM PMSF, 1x protease inhibitor (Sigma #p8340), and 1x phosphatase inhibitor cocktail (Sigma #p5726). Cells were intermittently incubated with the lysis buffer on ice and vortexed three times before being spun down for 10 minutes at 14,000 rpm. Protein concentrations were measured using bovine serum albumin for the protein standard and protein assay dye (Bio-Rad). A total of 40 μg of protein per condition was denatured with SDS-PAGE sample buffer and then loaded into polyacrylamide SDS-PAGE gels. The gels were run at 120 volts for 80 minutes and then transferred onto PVDF membranes (Millipore). The membranes were then blocked with 5% nonfat milk and probed with the appropriate primary antibody: SerpinB3 (Invitrogen PA5-30164, 1:5000), JAM-A (B&D Biosciences 612120, 1:1000), cathepsin L (ThermoFisher BMS1032, 1:5000) and c-Myc (Cell Signaling Technology 5605, 1:5000). β*-*Actin (Bio-Rad 12004163, 1:10,000) was used as a loading control. Secondary antibodies specific to the species of the primary antibody conjugated to horseradish peroxidase were added to the membranes: anti-rabbit (Invitrogen) and anti-mouse (EMD Millipore). Membranes were developed with Pierce ECL 2 Western Blotting Substrate (Thermo Scientific) onto film. For some experiments, secondary antibody was conjugated to StarBright 700 (Bio-Rad 12004161). For these experiments, a Bio-Rad Chemidoc MP was used to image the blots.

### Affinity purification of His-tagged JAM-A

T4121 CSCs were transiently transfected with N-terminal His-tagged full-length JAM-A (Sinobiologicals HG10198-NH). The His-tagged JAM-A was pulled down and isolated with nickel beads. Mass spectrometric analysis was used to identify binding partners that were pulled down along with JAM-A. For protein digestion, the bands were cut from the gel, washed/destained in 50% ethanol/5% acetic acid and then dehydrated in acetonitrile. The bands were then reduced with DTT and alkylated with iodoacetamide prior to in-gel digestion. Bands were digested overnight in-gel using trypsin. The peptides that were formed were extracted from the polyacrylamide in 50% acetonitrile with 5% formic acid. These extracts were combined and evaporated to <10 μL in a Speedvac and then resuspended in 1% acetic acid. The LC-MS system was a Finnigan LTQ-Obitrap Elite hybrid mass spectrometer system. The HPLC column was a Dionex 15 cm x 75 μm id Acclaim Pepmap C18, 2 μm, 100 Å reverse phase capillary chromatography column. The digest was analyzed using the data-dependent multitask capability of the instrument acquiring full-scan mass spectra to determine peptide molecular weights and product ion spectra to determine amino acid sequence in successive instrument scans. The data were analyzed by using all CID spectra collected in the experiment to search the human UniProtKB database with the search program Mascot. These partners were cross-referenced with the contaminant repository for affinity purification to remove negative controls.

### Immunostaining

Cells were plated onto coverslips in 6 well plates, fixed with 4% paraformaldehyde, blocked in goat serum with 0.1% Triton X-100, and then incubated with the appropriate primary antibody (SerpinB3 PA5-30164 Invitrogen, 1:500), JAM-A (Santa Cruz sc-53623, 1:500) followed by a species-specific secondary antibody. Secondary antibodies were as follows, goat-anti mouse Alexa Fluor 555 (ThermoFisher) and goat-anti rabbit Alexa Fluor 488 (ThermoFisher). The cells were then stained with Hoechst 33342 (Invitrogen H3570, 1:3000) before being mounted with Vectashield (Vector Labs) onto glass cover slides and imaged using a confocal microscope.

### Lysosomal membrane permeability post irradiation

T4121 tumor cells were treated with 5 Gy radiation. After 6 hours, 2 μg/ml acridine orange (Sigma A6014) was added for 30 minutes, and media was replaced before live cells were imaged.

For cathepsin L staining, tumor cells were treated with 5 Gy radiation and fixed with 4% formaldehyde. Cathepsin L antibody (ThermoFisher BMS1032, 1:1000) was added overnight, secondary antibody was then added (goat-anti mouse Alexa Fluor 488, ThermoFisher), and Hoechst 33342 (Invitrogen H3570, 1:3000) was utilized as a nuclear counterstain.

### Confocal microscopy

All images were taken with an inverted Leica SP8 confocal microscope at 40x magnification at room temperature. The LASX software was utilized for image acquisition. For image analysis, Fiji software was utilized. For quantification, images were split into individual channels, and the “integrated density” tool was utilized to quantify intensity of each channel per image. When quantifying the total number of cells, nuclei were counted manually.

### Immunohistochemistry on human glioblastoma

Standard immunohistochemistry analysis was performed on two patient specimens with a diagnosis of primary IDH-wild-type GBM using SerpinB3 antibody (Invitrogen PA5-30164) diluted at 1:1000. Four-micrometer thick sections of FFPE tissue on charged slides were baked in the oven at 60C for 60 minutes before being deparrafinized and re-hydrated. Antigen retrieval was achieved using a pH6 retrieval buffer (Biocare Reveal). Slides were cooled to room temperature and washed in TBS before neutralizing endogenous peroxidase (Biocare Peroxidase 1). Slides were then treated with a serum-free casein background block (Biocare Background Sniper) before pre-incubation in a 10% goat serum block for 60 minutes. Primary antibody was then added to the slides for overnight incubation at 4C. After incubation, slides were washed well with TBS-T before incubating in HRP polymer (Biocare MACH 4 Universal HRP Polymer). Finally, reaction products were visualized with DAB (Biocare Betazoid DAB Chromogen Kit). Slides were then counterstained with hematoxylin, dehydrated and mounted with xylene-based mounting media.

### Stable transduction with lentiviral shRNA and overexpression construct

MISSION® pLKO.1-puro Non-Mammalian shRNA Control Plasmid (SHC002) and SerpinB3 shRNA plasmids TRCN0000373440 (KD1), TRCN0000373501 (KD2), TRCN0000052398 (KD3) and TRCN0000373500 (KD4) were purchased from Sigma. Lentivirus was packaged in 293T cells using psPAX2 and pMD2G using calcium phosphate transfection, and media containing lentiviral particles were collected. This supernatant containing lentiviral particles was concentrated using PEGit virus precipitation solution according to the manufacturer’s protocol (System Biosciences). JAM-A knockdown constructs were as follows: TRCN0000061650 (KD1) and TRCN0000061649 (KD2).

Prior to transfection, CSCs were grown adherently on 6 well plates pretreated with Geltrex (Life Technologies). Lentivirus was added to and incubated with the cells for 24 hours. Then cells were grown in their appropriate media for 24 hours, after which selection with puromycin was initiated. Transfected cells were incubated in media with puromycin (1 mg/mL stock) at 1:333 for 48 hours. Stably transfected cells were maintained in their regular media plus puromycin at 1:4000.

### Cellular viability

Cellular viability was measured by plating each line of interest in triplicate in a 96-well plate at a density of 1000 cells/100 μL media per well. ATP levels at day 0 and day 7 were measured using CellTiter-Glo Luminescent Cell Viability Assay (Promega). For analysis, day 7 was normalized to the day 0 measurement.

To measure cell count over time, cells were plated in triplicate in a Geltrex-coated 96-well plate at a density of 1000 cells/100 μL media per well. The 96-well plate was then placed in the IncuCyte SX5 Live-Cell Analysis Instrument, and images were taken every 8 hours for 7 days. The cell-by-cell software was then utilized to determine cell count per well, and these values were normalized to the time 0 cell count.

### NanoString

RNA was isolated using an RNeasy mini kit (Qiagen), and then the nCounter® PanCancer Pathways Panel was used to analyze RNA expression. Two tumor models (T4121 and DI318) were analyzed in triplicate in each condition (non-target and SerpinB3 knockdown (KD2)). nSolver version 4.0 was utilized to determine pathway alterations.

### TGF-β ELISA

R&D systems human TGF-β1 DuoSet ELISA catalog# DY240 was used to quantify TGF-β1 in vitro from conditioned media isolated at day 2 after plating 200,000 cells per well in a 12 well plate with 1.5 ml of complete Neurobasal media. Output was normalized to total protein concentration in the pellet to control for changes in cell viability.

### Radiation treatment

A total of 50,000 cells per well were plated in triplicate in a 12 well plate. Cells were then irradiated with varying doses of radiation. On day 2, cell viability was measured using CellTiter-Glo Luminescent Cell Viability Assay (Promega). Viability was normalized to the untreated control for each condition and graphed as a percentage of the total.

### L-leucyl-L-leucine methyl ester

A total of 4,000 cells per well was plated in 96 well plate in quintuplets. L-leucyl-L-leucine methyl ester (LLME; Cayman #16008) was added over a range of concentrations. After 7 days of treatment, cell viability was measured, and half-maximal inhibitory concentrations (IC_50_) for each condition were calculated.

### Limiting-dilution analysis

Cells were plated at 100 cells per well in 12 wells of a 96 well plate, and two-fold serial dilutions were performed. Twelve wells of each cell dose were plated. Limiting dilution plots and stem-cell frequencies were calculated using ELDA analysis (http://bioinf.wehi.edu.au/software/elda/index.html; (Hu and Smyth, 2009)).

### Intracranial implantation

Intracranial tumor transplants were performed as described previously (Bayik et al., 2020). NSG mice were anesthetized with inhaled isoflurane for the duration of the procedure. A total of 20,000 T4121 CSCs infected with control or SerpinB3 shRNAs were suspended in 10 µl Neurobasal null medium and stereotactically implanted into the left hemisphere ∼2.5 mm deep into the brain. In relevant experiments, on day 10 after implantation, mice were anesthetized with xylazine (0.13 mg/mouse) and ketamine (1.3 mg/mouse) and exposed to 2 Gy radiation for either 3 or 5 days (PANTAK) starting 10 days post-tumor implantation and shielding the body with lead. Mice were monitored for neurologic signs and weight loss and deemed at endpoint when exhibiting any of these symptoms. Endpoint mice were transcardially perfused using 4% paraformaldehyde, and the brains were dissected for histological analysis after at least 48 hours in 4% paraformaldehyde. All experiments were performed in compliance with institutional guidelines and were approved by the Institutional Animal Care and Use Committee of the Cleveland Clinic (protocol 2019-2195 and 2019-2299).

### Cell death

For caspase activity assays, cells were plated in quintuplicate at 10,000 cells/well in 96 well plates for 48 hours. Caspase 3/7 activity was determined with the Caspase-Glo 3/7 assay (Promega) and caspase activity was normalized to cell number by performing the CellTiter-Glo Luminescent Cell Viability Assay on the duplicate plate.

Additionally, cells were plated for imaging in an IncuCyte as described above, and 5 μM Caspase 3/7 dye was included in the media (Sartorius #4704). The number of red nuclei (indicating active caspase 3/7) was divided by the area confluence per well. These values were then normalized to the non-target control.

For annexin V and propidium iodide assay, 25,000 cells/well were plated in 1.5 ml of Neurobasal media. After 48 hours, a single-cell suspension was obtained, and FITC-labeled annexin V and propidium iodide were added in accordance with the protocol (BioLegend #640914). Samples were run on an LSR Fortessa flow cytometer (BD Biosciences) with a minimum of 10,000 events collected. Single cells were gated, and the percentage of annexin V- and PI-positive cells was determined.

### Real-time reverse transcription polymerase chain reaction

RNA was collected from cells using an RNeasy kit (Qiagen). RNA concentrations were measured using a NanoDrop spectrophotometer, and cDNA was synthesized with qScript synthesis reagent (Quanta Biosciences). qPCR was run with the primers shown in Supplemental Table 2 using SYBR-Green Mastermix (SA Biosciences) and an Applied Biosystems QuantStudio 3. During analysis, threshold cycle numbers were normalized to GAPDH levels.

### Depmap RNAi

The RNAi DEMETER2 analysis framework was utilized to determine the gene dependency of SerpinB3 (McFarland et al., 2018). Data was accessed from https://depmap.org/R2-D2/ on 12/12/2021 and relied on three large RNAi datasets (Marcotte et al., 2016; McDonald et al., 2017; Tsherniak et al., 2017).

### Statistical analysis

For two-group comparisons, p-values were calculated using ANOVA. For multiple group comparisons, one-way ANOVA with Dunnett’s multiple comparisons test was used as indicated in the figure legends. Log-rank tests were used for survival analysis. GraphPad Prism 6 was used for statistical tests. All in vitro experiments were done in at least technical triplicates for each experimental group, and multiple independent experiments were performed. The Grubb’s test was performed to determine whether any outliers were statistically different. Statistical details can be found in figure legends. p<0.05 was considered statistically significant. *, p< 0.05; **, p< 0.01; ***, p< 0.001.

## Author Contributions

Conceptualization: AL, SM, JV, JDL; Data Analysis: AL, ST, JV, DB, DJS, KM, EEM-H, KD, MM, JH, KM, AS Project Administration: AL, JDL; Supervision: UN, SB, CMH, YU, JDL; Funding acquisition: AL, CMH, JDL; Writing – original draft: AL, EEM-H, JDL; Writing – reviewing & editing: All authors

## Acknowledgements

We thank Drs. William Schiemann, Mark Jackson, and Alex Huang (Case Western Reserve University) for critical feedback and Gary Silverman (Washington University in St. Louis School of Medicine) for discussions and advice about lysosomal assessments. We thank Ms. Karen Kasler and Dr. Robert Fairchild for their assistance with Nanostring analysis. We thank Belinda Willard for assistance with the mass-spec and proteomics analysis. We also thank members of the Lathia laboratory including Katie Troike, Samuel Sprowls, Salma Ben Salem, Kristen Kay, Juyeun Lee, and Sabrina Wang for insightful discussions, and Sadie Johnson for their assistance with mouse work. We thank Ms. Amanda Mendelsohn for illustration assistance.

## Supplemental Figure Legends

**Supplemental Figure 1**

A) SerpinB3 expression was measured via western blot in T4121 tumor cells containing JAM-A shRNA. B) SerpinB3 and JAM-A expression in four human GBM tumor samples. C) Immunohistochemistry of SerpinB3 in two IDH-wild-type glioblastoma patients. D) IncuCyte growth data, with cell count every 8 hours for 7 days, graphed as fold change relative to time 0. E) Western blot of SerpinB3 knockdown 3 (KD3) and knockdown 4 (KD4) F) Cell viability was measured with CellTiter Glo at day 7 in multiple GBM PDX models with KD3 and KD4 shRNA constructs (five technical replicates per condition, per tumor model). G) SerpinB3 dependency of 31 GBM cells lines by RNAi screen. p<0.05 was considered statistically significant. *, p<0.05; **, p<0.01; ***, p<0.001 as determined by one-way ANOVA with Dunnett’s multiple comparisons. Error bars represent standard deviation.

**Supplemental Figure 2**

A) Overall unbiased clustering from the NanoString platform analysis of DI318 and T4121 PDX models with non-target and SerpinB3 KD2 shRNA. B-D) Volcano plot comparing NT control to SerpinB3 knockdown, with members of each pathway highlighted in yellow. E) mRNA expression of Notch2 and Jagged2 after SerpinB3 knockdown. *, p<0.05; **, p<0.01; ***, p<0.001 as determined by one-way ANOVA with Dunnett’s multiple comparisons. The highest solid black bar represents p< 0.01.

**Supplemental Figure 3**

A) Caspase 3/7 activity in T387 cells depleted for SerpinB3 with shRNA KD3 and KD4. p<0.05 was considered statistically significant (five technical replicates per condition, per tumor model). B) Six hours post irradiation, T387 cells were fixed with paraformaldehyde and stained for cathepsin L. Images from nine random fields were taken for each group. The integrated density of cathepsin L per cell was determined and compared between the irradiated and control groups. C) Western blot demonstrating total expression of SerpinB3 and cathepsin L 6 hours after irradiation. *, p<0.05; **, p<0.01; ***, p<0.001 as determined by one-way ANOVA with Dunnett’s multiple comparisons. Error bars represent standard deviation.

**Supplemental Figure 4**

A) Schematic of in vivo radiation experiment with 10 Gy total. B-G) A total of 20,000 tumor cells per condition were intracranially injected into mice. Ten days after irradiation, mice heads received 2 Gy of radiation every other day for 3 days (total of 6 Gy) to the head. B) All treatment groups plotted with median survival values. The groups were subsequently divided into: C) Non-target with or without radiation, D) SerpinB3 knockdown 1 with or without radiation, E) SerpinB3 knockdown 2 with or without radiation. F) SerpinB3 dependency across three large RNAi screens on tumor cell lines from a variety of tumors. p<0.05 was considered statistically significant. *, p<0.05; **, p<0.01; ***, p<0.001 as determined by log-rank test.

