## Supplementary figures and images for "SerpinB3 drives cancer stem cell survival in glioblastoma"

### Supplemental Figure 1

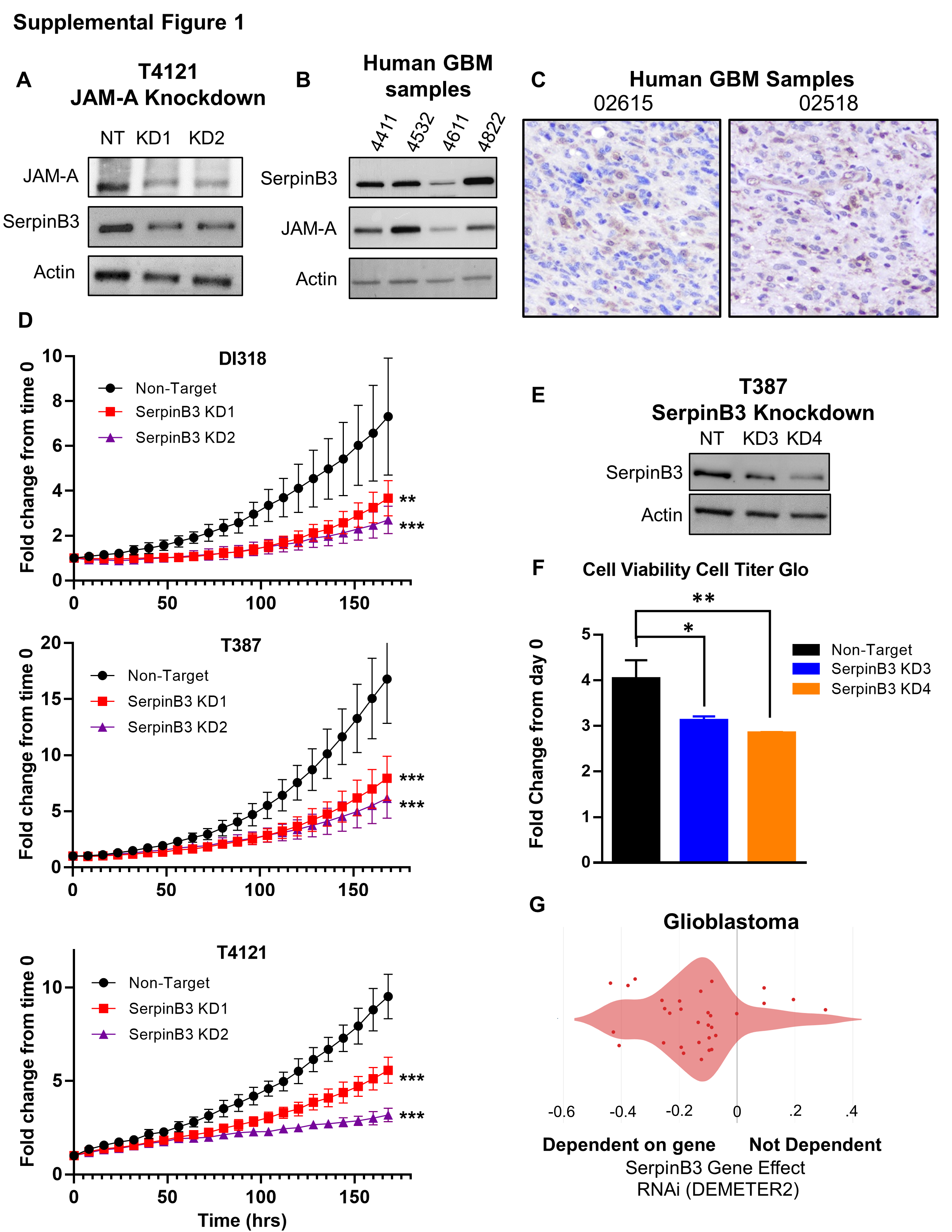

### Supplemental Figure 2

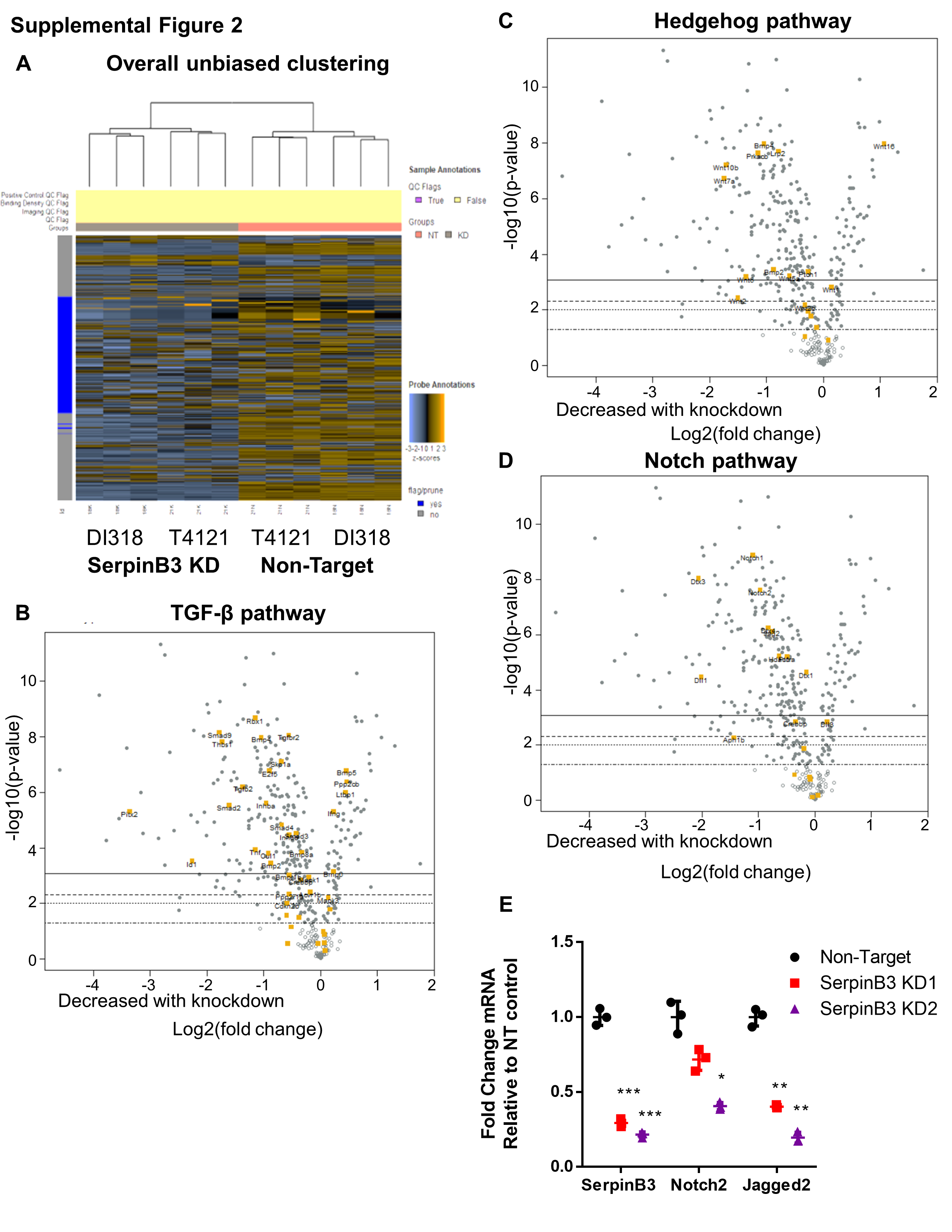

### Supplemental Figure 3

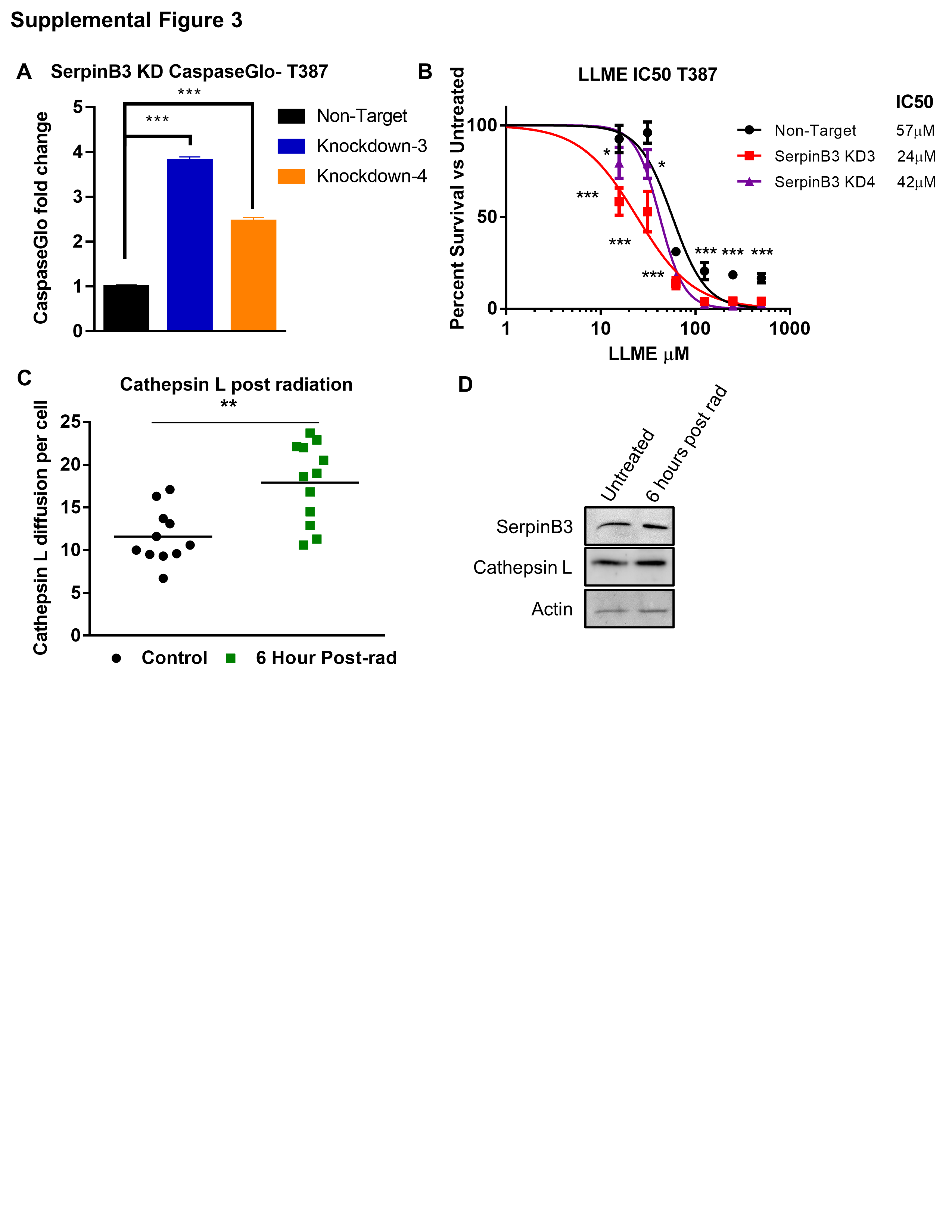

### Supplemental Figure 4

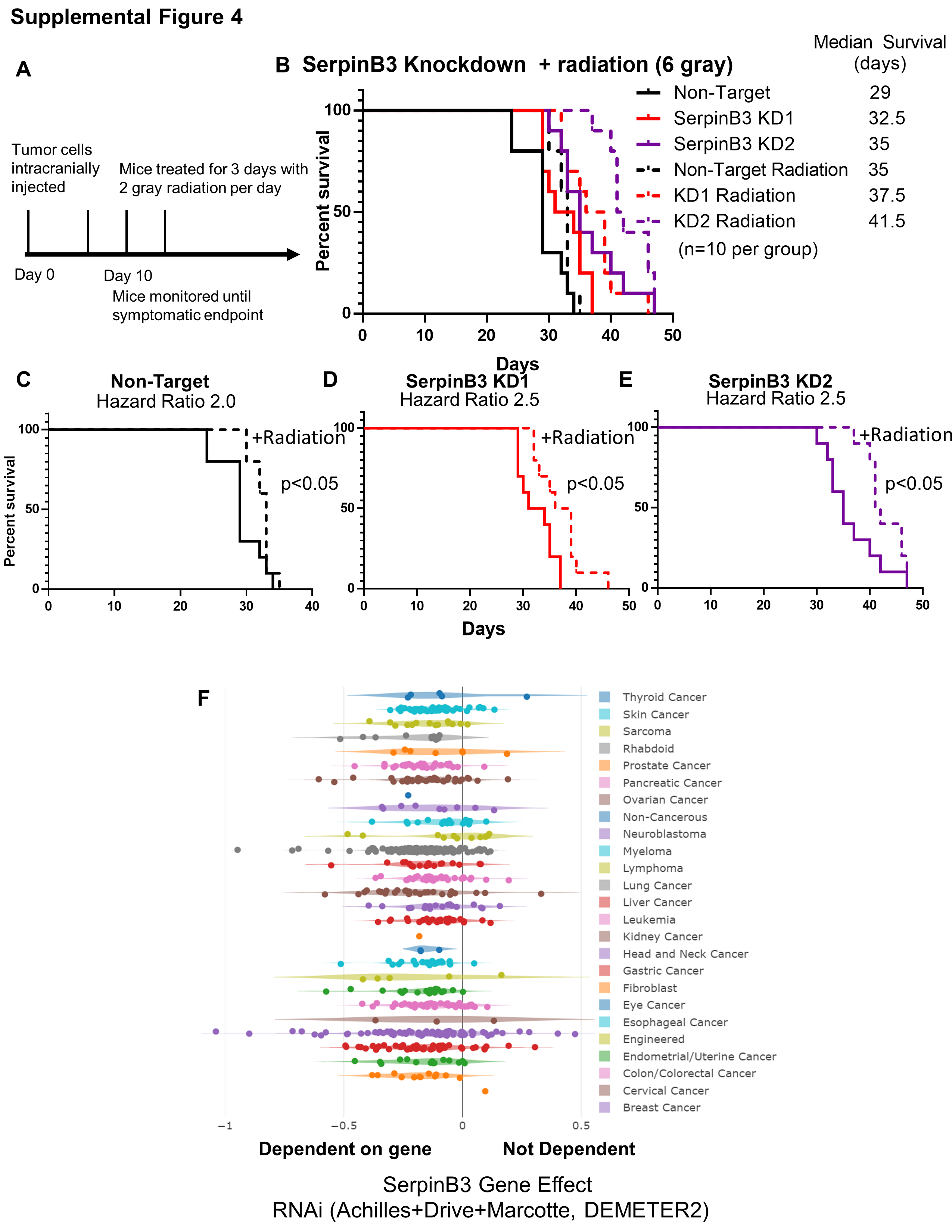

### Supplemental Tables 1 and 2

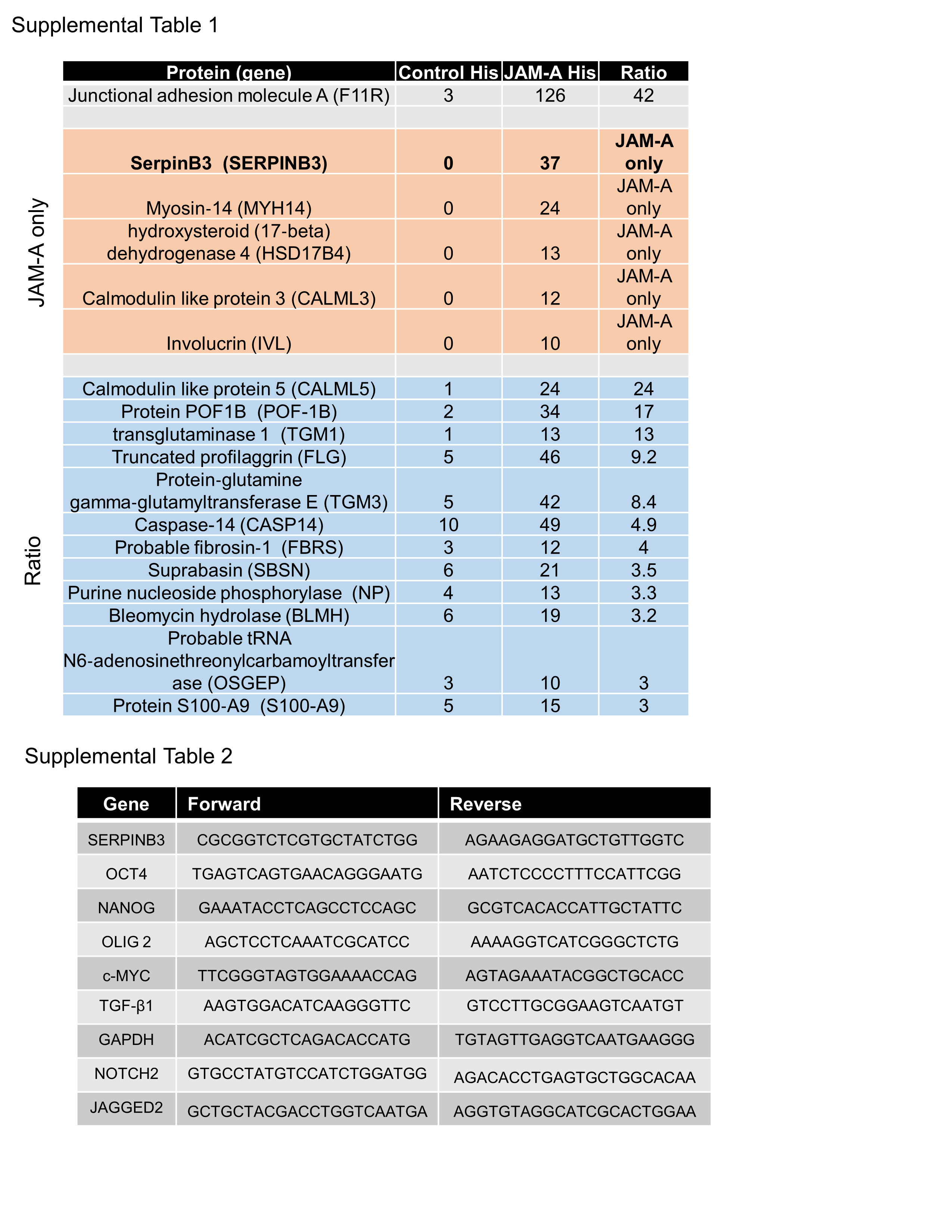
